# A dynamic model of the ABA Signaling pathway with its core components: translation rate of PP2C determines the kinetics of ABA-induced gene expression

**DOI:** 10.1101/2021.12.08.471820

**Authors:** Ruth Ndathe, Renee Dale, Naohiro Kato

## Abstract

The abscisic acid (ABA) signaling pathway is the key defense mechanism against drought stress in plants, yet the connectivity of cellular molecules related to gene expression in response to ABA is little understood. A dynamic model of the core components of the ABA signaling pathway was built using ordinary differential equations to understand the connectivity. Parameter values of protein-protein interactions and enzymatic reactions in the model were implemented from the data obtained by previously conducted experiments. On the other hand, parameter values of gene expression and translation were determined by comparing the kinetics of gene expression in the model to those of ABA-induced *RD29A* (response to desiccation 29A) in actual plants. Based on the analyses of the optimized model, we hypothesized that the translation rate of PP2C (protein phosphatase type 2C) is downregulated by ABA to increase the ABRE (ABA-responsive element) promoter activity. The hypotheses were preliminarily supported by newly conducted experiments using transgenic Arabidopsis plants that carry a luciferase expression cassette driven by the RD29A promoter (*RD29A::LUC*). The model suggests that identifying a mechanism that alters PP2C translation rate would be one of the next research frontiers in the ABA signaling pathway.

## Introduction

Plants possess defense mechanisms against drought (Basu *et al*., 2016; Kumar *et al*., 2018; Takahashi *et al*., 2020a). One of the major mechanisms is the abscisic acid (ABA) signaling pathway. ABA is a phytohormone that is produced under the drought stress conditions (Zeevaart & Creelman, 1988; Sauter *et al*., 2001; Ikegami *et al*., 2008). The ABA signaling pathway has been well-characterized, leading to downstream ABA responses such as stomatal closure and gene expression that help the plant to acquire drought stress resistance (Steuer *et al*., 1988; Fujii *et al*., 2009; Umezawa *et al*., 2009). The most upstream of the core components in the ABA signaling pathway is ABA-receptors named pyrabactin resistance/pyr1-like/ regulatory components of ABA receptors (PYR/PYL/RCAR) that bind ABA and in turn interact with different protein phosphatase 2Cs (PP2Cs), namely aba insensitive1/2 (ABI1/ABI2), hypersensitive to aba1/2 (HAB1/HAB2), aba-hypersensitive germination 3 (AHG3/PP2CA), and highly aba induced 1/2/3 (HA1/2/3). Without the PYR interaction, these PP2Cs inhibit SNF1-related protein kinase 2s (SnRK2s) that include SnRK2.2, SnRK2.3 and SnRK2.6. (Rodriguez *et al*., 1998; Gosti *et al*., 1999; Merlot *et al*., 2001; Saez *et al*., 2004; Ma *et al*., 2009; Melcher *et al*., 2009; Nishimura *et al*., 2009; Park *et al*., 2009; Santiago *et al*., 2009; Yin *et al*., 2009; Soon *et al*., 2012). Activated SnRK2s phosphorylate ABA-responsive elements (ABRE) binding factors 1/2/3/4 (ABF1/2/3/4). These phosphorylated transcription factors bind ABREs on a regulatory region of ABA-induced genes (Choi *et al*., 2000; Uno *et al*., 2000; Yoshida *et al*., 2015). Alternatively, the activated SnRK2, namely SnRK2.6 kinase, phosphorylate the slow-anion channels (SLAC1) leading to their activation and subsequently lead to stomatal closure due to anion and K^+^ efflux and eventual solute loss from the guard cells (Schroeder *et al*., 1984; Geiger *et al*., 2009; Lee *et al*., 2009; Albert *et al*., 2017).

The ABA signaling pathway has been mathematically modeled to help understand the ABA signaling pathway in guard cells leading to stomatal closure (Li *et al*., 2006; Albert *et al*., 2017; Maheshwari *et al*., 2019; Maheshwari *et al*., 2020). These works have led to the determination of new predictions and hypotheses in the ABA signaling pathway, for example, the role of feedback regulation, ROS, Ca^2+^, pH, and heterotrimeric G-protein signaling in ABA-induced stomatal closure (Li *et al*., 2006; Albert *et al*., 2017; Maheshwari *et al*., 2019). In addition, the additive effect of ABA and salt stress on ABA and drought-responsive expression of genes was also explained using mathematical modeling (Lee *et al*., 2016).

The ABA signaling pathway has additional regulatory mechanisms, which are feedback and post-translational regulations. The feedback regulation involves upregulation of PP2C genes, which eventually results in enhanced deactivation of SnRK2s (Rodriguez *et al*., 1998; Saez *et al*., 2004; Fujita *et al*., 2009; Wang *et al*., 2019). It also includes the upregulation of ABF genes, which increases ABF expression (Wang *et al*., 2019). These regulatory elements are thought to affect gene expression kinetics. The post-translation regulation involves phosphorylation of PYL by the target of rapamycin (TOR) protein kinase (Wang *et al*., 2018). On the other hand, Raptor, the TOR associated protein, is phosphorylated by SnRK2s, leading to TOR kinase inhibition (Wang *et al*., 2018). In another study, TOR was found to suppress ABA-responses by phosphorylating *Arabidopsis thaliana* yet another kinase (AtYAK1) (Forzani *et al*., 2019) that is a positive regulator of ABA-mediated signal responses (Kim *et al*., 2016). Therefore, TOR was proposed to be a post-translation regulator in the ABA signaling pathway. E3-ligases are another post-translational regulator which promotes the degradation of ABA signaling components, including PP2CA (Wu *et al*., 2016), SnRK2.6 (Ali *et al*., 2019), and PYL5/7/8/9 (Zhao *et al*., 2017).

Network connectivity of these additional regulatory mechanisms to the core components is little understood. Dynamic modelling can allow us to better understand their role in the ABA signaling pathway. Dynamic modelling is a powerful tool that integrates extensive experimental data of pathway components, improving our understanding of the signaling pathway dynamics and making novel hypotheses and predictions (Poolman *et al*., 2004; Aldridge *et al*., 2006; Janes & Yaffe, 2006; Thakar *et al*., 2007). *In vitro* parameters for many of the interactions of the core components in the ABA signaling pathway have been experimentally determined, allowing us to create a dynamic model.

The purpose of this study is to build a dynamic model consisting of the core components with fixed parameter values that were previously obtained by experiments. Approximate curve fitting of the model output to actual plant data was conducted by optimizing parameter values of transcription and translation, which were not determined previously. In this report, we describe how we built, optimized, and validated the model. The resulting model suggested two novel hypotheses, which were supported by preliminary experiments. This model can be expanded to investigate the roles of additional regulatory mechanisms in future studies.

## Description

### Construction of the dynamic model

A previous study defined a minimal set of core components that activate the ABFs, leading to ABA-induced gene expression in the ABA signaling pathway (Fujii *et al*., 2009). The components are ABA, PYR/PYL/RCAR, PP2Cs (ABI1/2 and HAB1/2), SnRK2s (SnRK2.2/3/6), ABFs (ABF2/3/4), and ABRE. Other studies have determined that the PP2CA phosphatases dephosphorylate phosphorylated ABFs (Antoni *et al*., 2012; Lynch *et al*., 2012). In addition, another study identified MAP3K phosphorylates SnRK2s (Takahashi *et al*., 2020b). These two reactions were included in the model. We also included the feedback regulation in which the expression of PP2C, PP2CA, and ABF genes are upregulated by the ABRE promoter activity (Wang *et al*., 2019). A set of 21 ordinary differential equations representing biochemical reactions of each component were constructed based on the law of mass action (Fig. **1**). Homologous proteins with redundant function are modeled as a single protein. Initial values of variables and values of parameters in the equations were obtained from the literature (Table **1**). The equations, initial conditions (concentrations), and parameter values were then compiled and numerically analyzed with MATLAB R2020b SimBiology (MathWorks) with default settings.

**Figure 1.**
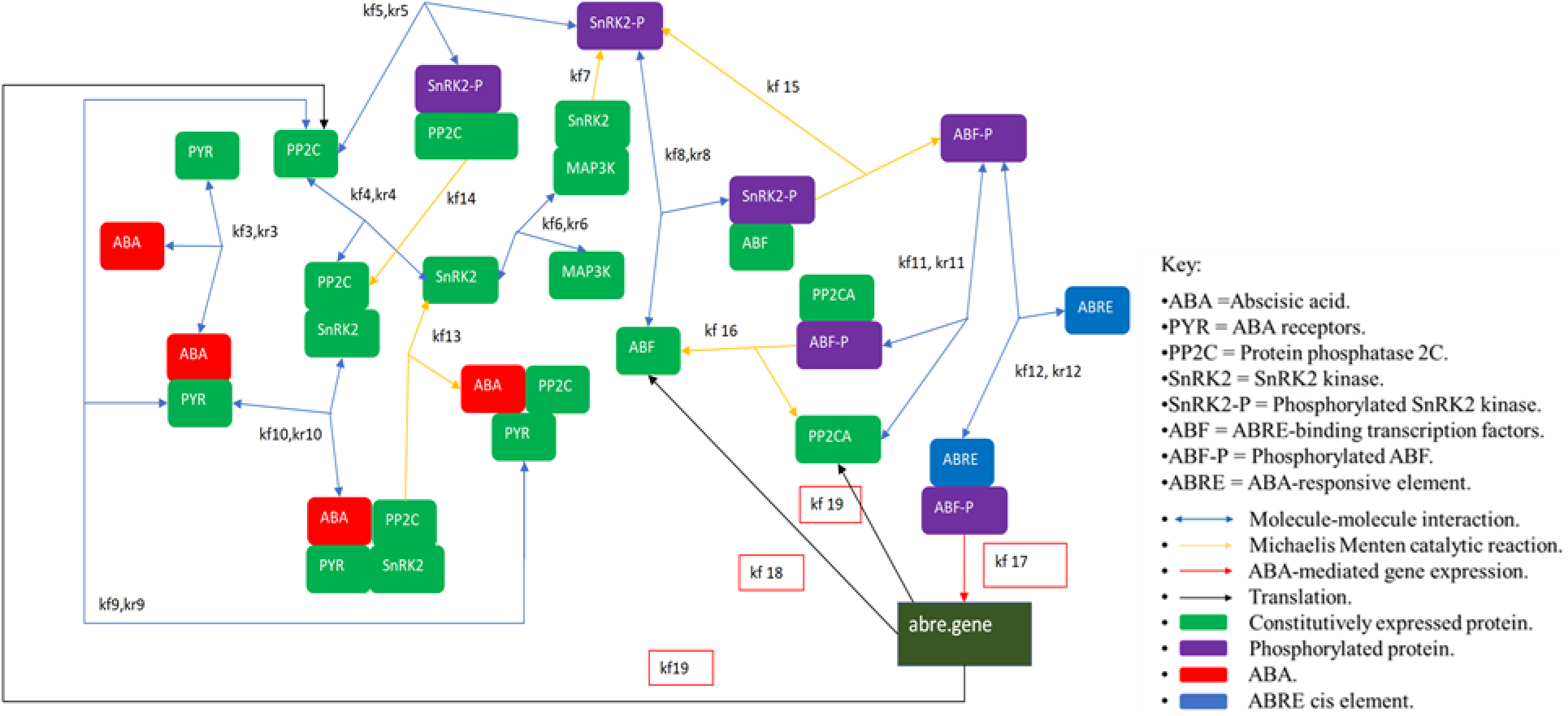
A schematic mass-action model of the ABA-signaling pathway with its core components. Rectangles and arrows represent variables and reactions, respectively. Identifiers of parameters in each reaction are shown as kf or kr with unique number. Parameters optimized in this study are indicated with a red frame. The values of each parameter are shown in Table **1**.

**Table 1.** Curated values from literature and the values chosen as parameters for the model. Each reaction in the model was shown with the respective parameter and the source from which the value was obtained.

| Description. | Reference. | Value found in the literature. | Parameter name in the model. | Value used in the model. | Fixed in the model*. |
| --- | --- | --- | --- | --- | --- |
| Transcription of constitutively expressed genes | (Hausser *et al.*, 2019) | < translation rate | kf1 | 1 hr<sup>-1</sup> | ✓ |
| Translation of constitutively expressed genes | (Hausser *et al.*, 2019) | < 10,000 hr<sup>-1</sup> | kf2 | 4.5 hr<sup>-1</sup> | ✓ |
| ABA and PYR binding | (Dupeux *et al.*, 2011) | $K_D = 65 \mu\text{M}$ | kf3<br>kr3 | 1000 $\mu\text{M}^{-1} \text{s}^{-1}$<br>65000 $\text{s}^{-1}$ | ✓ |
| PP2C and SnRK2 binding | (Soon *et al.*, 2012) | $\text{IC}_{50}$<br>2 $\mu\text{M}$ – 8 $\mu\text{M}$ | kf4<br>kr4 | 1000 $\mu\text{M}^{-1} \text{s}^{-1}$<br>0.1 $\text{s}^{-1}$ | ✓ |
| PP2C and SnRK2-P binding | (Xie *et al.*, 2012) | $K_M = 0.097 \mu\text{M}$ | kf5<br>kr5 | 1000 $\mu\text{M}^{-1} \text{s}^{-1}$<br>97 $\text{s}^{-1}$ | ✓ |
| SnRK2 and MAP3K binding | (Ghose, 2019) | $K_M = 23 \mu\text{M}$ | kf6<br>kr6 | 1000 $\mu\text{M}^{-1} \text{s}^{-1}$<br>23000 $\text{s}^{-1}$ | ✓ |
| Phosphorylation of SnRK2 by MAP3K | (Ghose, 2019) | $k_{cat} = 14 \text{ s}^{-1}$ | kf7 | 14 $\text{s}^{-1}$ | ✓ |
| SnRK2-P and ABF binding | (Xie *et al.*, 2012) | $K_M = 19.3 \mu\text{M}$ | kf8<br>kr8 | 1000 $\mu\text{M}^{-1} \text{s}^{-1}$<br>19300 $\text{s}^{-1}$ | ✓ |
| PYR.ABA and PP2C binding | (Dupeux *et al.*, 2011) | $K_D = 30 \text{ nM}$ | kf9<br>kr9 | 1000 $\mu\text{M}^{-1} \text{s}^{-1}$<br>30 $\text{s}^{-1}$ | ✓ |
| PYR.ABA and PP2C.SnRK2 binding | (Dupeux *et al.*, 2011) | $K_D = 30 \text{ nM}$ | kf10<br>kr10 | 1000 $\mu\text{M}^{-1} \text{s}^{-1}$<br>30 $\text{s}^{-1}$ | ✓ |
| ABF-P and PP2CA binding | (Pan *et al.*, 2015) | $K_M = 11.15 \mu\text{M}$ | kf11<br>kr11 | 1000 $\mu\text{M}^{-1} \text{s}^{-1}$<br>11150 $\text{s}^{-1}$ | ✓ |
| ABF-P and ABRE binding | (Geertz *et al.*, 2012) | $K_D$ of DNA-protein binding<br>2 nM - 2 $\mu\text{M}$ | kf12<br>kr12 | 1000 $\mu\text{M}^{-1} \text{s}^{-1}$<br>2 $\text{s}^{-1}$ | ✓ |
| Release of SnRK2 from ABA.PYR.PP2C.SnRK2 complex. | (Bar-Even *et al.*, 2011) | Average $k_{cat}$ of enzyme reaction<br>10 $\text{s}^{-1}$ | kf13 | 10 $\text{s}^{-1}$ | ✓ |
| Dephosphorylation of SnRK2-P | (Xie *et al.*, 2012) | $k_{cat} = 0.924 \text{ s}^{-1}$ | kf14 | 0.924 $\text{s}^{-1}$ | ✓ |
| Phosphorylation of ABF by SnRK2-P | (Xie *et al.*, 2012) | $k_{cat} = 0.04 \text{ s}^{-1}$ | kf15 | 0.04 $\text{s}^{-1}$ | ✓ |
| Dephosphorylation of ABF-P by PP2CA | (Pan *et al.*, 2015) | $k_{cat} = 1.04 \text{ s}^{-1}$ | kf16 | 1.04 $\text{s}^{-1}$ | ✓ |
| Transcription of ABA induced genes | (Hausser *et al.*, 2019) | < translation rate | kf17 | 10 hr<sup>-1</sup> |  |
| Translation of feed-backed ABF | (Hausser *et al.*, 2019) | < 10,000 hr<sup>-1</sup> | kf18 | 200 hr<sup>-1</sup> |  |
| Translation of feed-backed PP2C and PP2CA | (Hausser *et al.*, 2019) | < 10,000 hr<sup>-1</sup> | Kf19 | 200 hr<sup>-1</sup> |  |
| Degradation of mRNA | (Hausser *et al.*, 2019) | mRNA degradation in HEK293 cells<br>0.06 hr<sup>-1</sup> | kf20, kf21 | 0.06 hr<sup>-1</sup> | ✓ |
| Degradation of protein | (Hausser *et al.*, 2019) | Protein decay rate in Hela cells<br>0.05 hr<sup>-1</sup> | kf22 to kf38 | 0.05 hr<sup>-1</sup> | ✓ |

\*Fixed in the model: ✓ indicates the value used in the model was not altered during model optimization

In the model, we assumed:

- ABA signal transduction occurs through molecule-molecular interactions; where the molecule could be a protein, a hormone, or DNA.
- Enzymatic reactions follow Michaelis-Menten kinetics.
- All molecules freely diffuse in the cell.
- The cell volume is 50 μm^3^.
- The Michaelis constant is 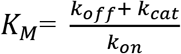 where *k_off_* is the dissociation rate constant, *k_cat_* is the catalytic rate constant, and *k_on_* is the association rate constant.
- A molecule associates with another molecule at a rate constant of, *k*_on_ = 1000 μM^−1^s^−1^ (Milo & Phillips, 2015).
- Proteins are generated by reactions of gene expression and protein translation, then subject to degradation.
- The concentration of a protein in a cell remains at 0.1 μM at a steady state without ABA activation and feedback regulation.
- A gene (mRNA) is expressed from a pair of gene loci that have a constitutively active promoter, then subjected to degradation.
- A gene (mRNA) that is expressed by a feedback regulation has an additional regulatory element (ABRE) in the same gene loci that have a constitutively active promoter.

In numerical analysis, the model was first run for 300 equivalent hours with the variable ABA (representing intracellular ABA) set at 0 μM. This allows the system to reach a quasi-steady state. After the 300 equivalent hours, the variable ABA was set to 100 μM. Changes of all variables in the model from the quasi-steady state was then monitored for another 300 equivalent hours. In this report, the time point when the variable ABA is changed is presented as time zero.

### Optimization of parameters, validation of the model, and analyzing identifiability of model parameters

To optimize selected model parameters, we approximately curve fit model output to experimental data. We focused on changes in the variable abre.gene, representing accumulated mRNA expressed from the ABRE promoter. Three parameters, 1. transcription of ABA-induced genes, 2. translation of feed-backed ABF, 3. translation of feedbacked PP2C and PP2CA, were manually changed to obtain qualitatively good fits to experimental data. The remaining model parameters were unchanged (fixed). To validate the model, we quantitatively evaluated changes of the variable abre.gene. Fold changes calculated by the model were compared to data previously published or data newly obtained in this study. To analyze identifiability on the dynamics of the variable abre.gene, we conducted sensitivity analysis using Calculate Sensitivity in Model Analyzer in SimBiology with default settings.

## Results

### Parameter values were obtained by literature curation

We curated previously published data to define parameters in the model of the ABA signaling pathway that activates the ABF, resulting in the activation of the gene promoter containing ABRE cis element. The summary of our curation is shown below (Table **1**).

While parameter values for protein-protein interactions and enzymatic reactions were characterized in *in vitro* studies using recombinant proteins, no studies related to parameter values of DNA-protein binding, gene expression, protein translation and degradation were found for the ABA signaling pathway. To this end, we implemented parameter values from studies using non-plant eukaryotic organisms. These parameters have a wide range to select from: 1. equilibrium dissociation constant between ABF-P (phosphorylated ABF) and the ABRE promoter (from 2 nM to 2 μM) (Geertz *et al*., 2012), 2. translation rate of protein from mRNA expressed by the ABRE promoter (less than 10,000 hr^−1^) (Hausser *et al*., 2019), 3. transcription rate of the ABRE promoter (slower than the translation rate) (Hausser *et al*., 2019). We selected the values of translation and transcription rates for genes at 4.5 hr^−1^ and 1 hr^−1^, respectively, and 2nM for (ABF-P)-(ABRE) binding. This is because an average rate of gene transcription in multicellular eukaryotes is 1 hr^−1^ (Hausser *et al*., 2019) while an average concentration of proteins involved in a signal transduction is 0.1 μM (Milo & Phillips, 2015). Setting translation rate at 4.5 hr^−1^ and transcription rate at 1 hr^−1^ makes the concentration of a protein at quasi-steady state to 0.1 μM without ABA and feedback regulation in our model. The affinity of (ABF-P)-(ABRE) binding was set at 2 nM to curve-fit kinetics of the variable abre.gene with actual gene expression (Fig. **2**). Protein degradation was set at 0.05 hr^−1^ (Hausser *et al*., 2019). Equilibrium dissociation constant between SnRK2 (non-phosphorylated SnRK2) and PP2C was set at 100 pM, representing complete inhibition of SnRK2 kinase activity by PP2C at an equal molar concentration (Soon *et al*., 2012).

**Figure 2.**
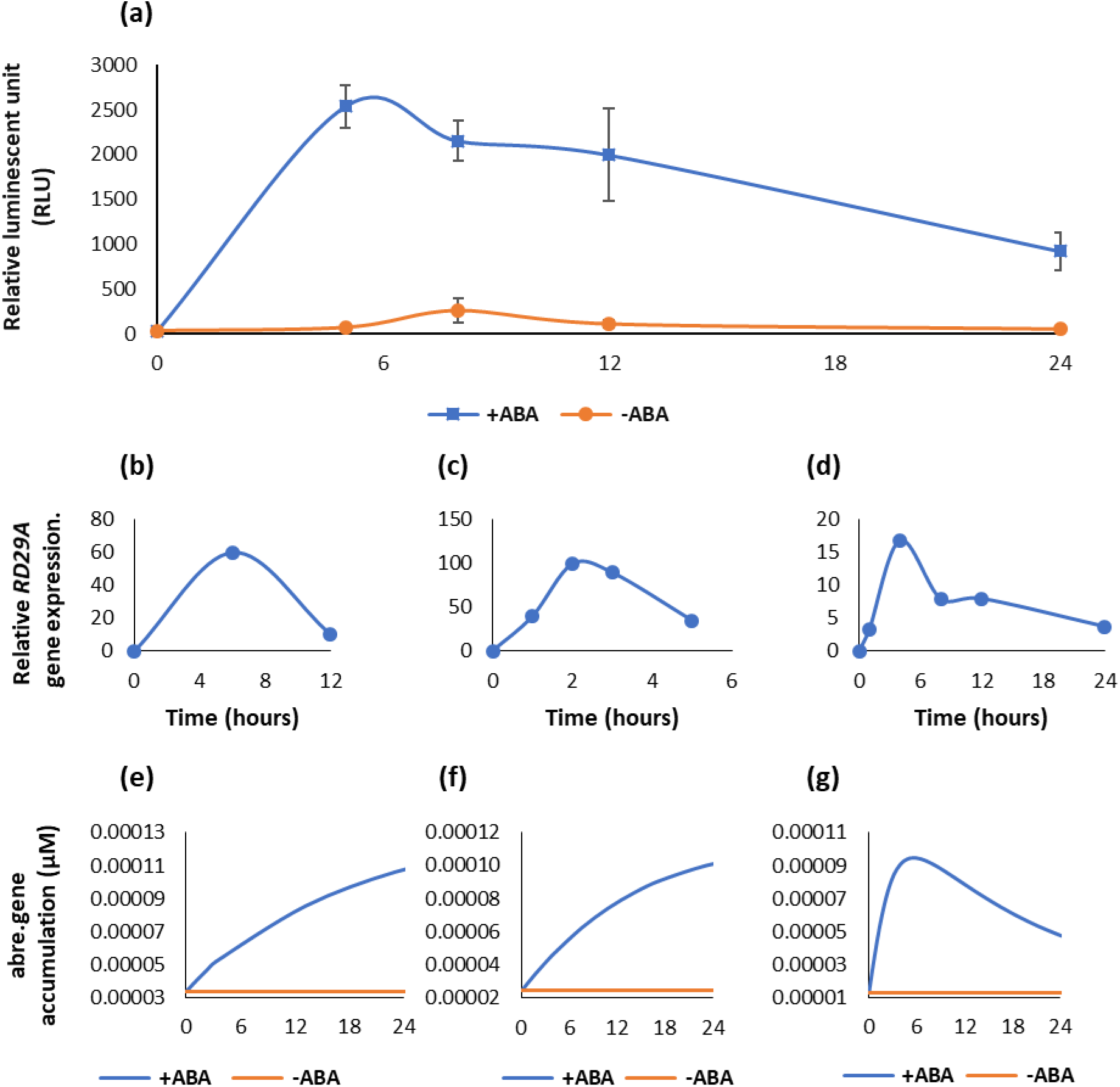
Dynamic model agrees with ABA-induced gene expression in real plants after optimization. (**a**) Kinetics of luciferase activity in the *RD29A::LUC* plant after exposing to 200μM ABA (+ABA) or DMSO for control (-ABA). The graph shows a mean of three independent experiments. Error bars represent standard error from the mean. (**b**) Kinetics of *RD29A* gene accumulation in the previously published data with 50 μM ABA in rice (Singh *et al*., 2015). (**c**) Kinetics of *RD29A* gene accumulation in the previously published data with 100 μM ABA in Arabidopsis (Lee *et al*., 2016). (**d**) Kinetics of *RD29A* gene accumulation in the previously published data with 10 μM ABA in Arabidopsis (Song *et al*., 2016). (**e**) Model output without feedback regulation (kf17 = 1 hr^−1^). (**f**) Model output with feedback regulation (adding reactions kf18= 4.5 hr^−1^ and kf19= 4.5 hr^−1^). (**g**) Model output with feedback regulation and optimized parameters (kf17=10 hr^−1^, kf18 = 200 hr^−1^, kf19 = 200 hr^−1^).

### The transcription rate of genes expressed by the ABRE promoter and the translation rate of feedback loop components ABF, PP2C, and PP2CA were optimized in the model to capture observed dynamics in experimental data

To understand the connectivity of the components, we compared the kinetics of gene expression in the model and experimental data in actual plants. Namely, we compared the simulation data of the variable abre.gene, which represents the accumulation of genes expressed by the ABRE promoter, to four independent data sets that were experimentally obtained using actual plants. One set of data was obtained by our new experiments using transgenic *Arabidopsis thaliana*. The transgenic plants carry the *RD29A::LUC* gene expression cassette that has been used to study the activity of the ABRE promoter (Zhan *et al*., 2012). The activity of ABRE promoter can be monitored by luminescence in near real-time in plants. The other three sets were obtained from previously published data that show a change in *RD29A* gene expressed from the native ABRE promoter in the genome of either *Arabidopsis thaliana* (Lee *et al*., 2016; Song *et al*., 2016) or *Oryza sativa* (rice) (Singh *et al*., 2015). Kinetics of the gene expression in the plants and the variable abre.gene were compared within the first 24 hours (Fig. **2**).

Experimental data from the transgenic *RD29A::LUC* plants showed transient activation of the ABRE promoter with an initial increase and then a decrease after 5 hours (Fig. **2a**). Similar transient expression of the *RD29A* gene were observed in non-transgenic plants, Arabidopsis and rice (Fig. **2b**, **c**, **d**) (Singh *et al*., 2015; Lee *et al*., 2016; Song *et al*., 2016). When we simulated kinetics of the variable abre.gene in the model without the feedback regulation on ABF, PP2C, and PP2CA (parameters kf18 and kf19), the kinetics were logarithmic upon adding ABA (Fig. **2e**). Addition of the feedback regulation had minor impact on the kinetics (Fig. **2f**). We then optimized the parameters so that kinetics of the gene expression in the model qualitatively agree with that in actual plants (Fig. **2g**). We namely altered the three parameters, the transcription rate constant of the ABRE promoter (parameter kf17) and the translation rate constants of ABF and PP2Cs (parameter kf18 and kf19, respectively) (Fig. **1** & Table **1**). These three parameters had not been determined previously, and studies in other eukaryotic cells indicate wide ranges of reasonable values (Table **1**). Hence, we selected the values within the ranges that made the kinetics of the variable abre.gene best fit to the actual plant data. The values kf17= 10 hr^−1^, kf18= 200 hr^−1^, and kf19=200 hr^−1^ fitted the kinetic curve with the actual plant reasonably (Fig. **2a**, **g**).

### Approximation of the model was validated by determining model responses to different doses of ABA or a set of gene null-mutations

To validate the model, we first compared the ABA-dose-dependent response in actual plants to the dynamics of the variable abre.gene (Fig. **3**). In the model, changes of the variable abre.gene increased in an ABA-dose dependent manner in the range from 0 to 200 μM (Fig. **3a**). With the *RD29A::LUC* transgenic plants, changes of luminescence increased in an ABA-dose dependent manner in the range from 0 to 200 μM (Fig. **3b**). This suggested that the model is approximated to actual plants with respect to ABA sensitivity although the response in the model seems to have narrower sensitivity against the ABA concentration (i.e., from 0 to 50 μM) compared to that in the actual plants (i.e., from 0 to 200 μM) (Fig. **3b**) (Gampala *et al*., 2001; Lee *et al*., 2016).

**Figure 3.**
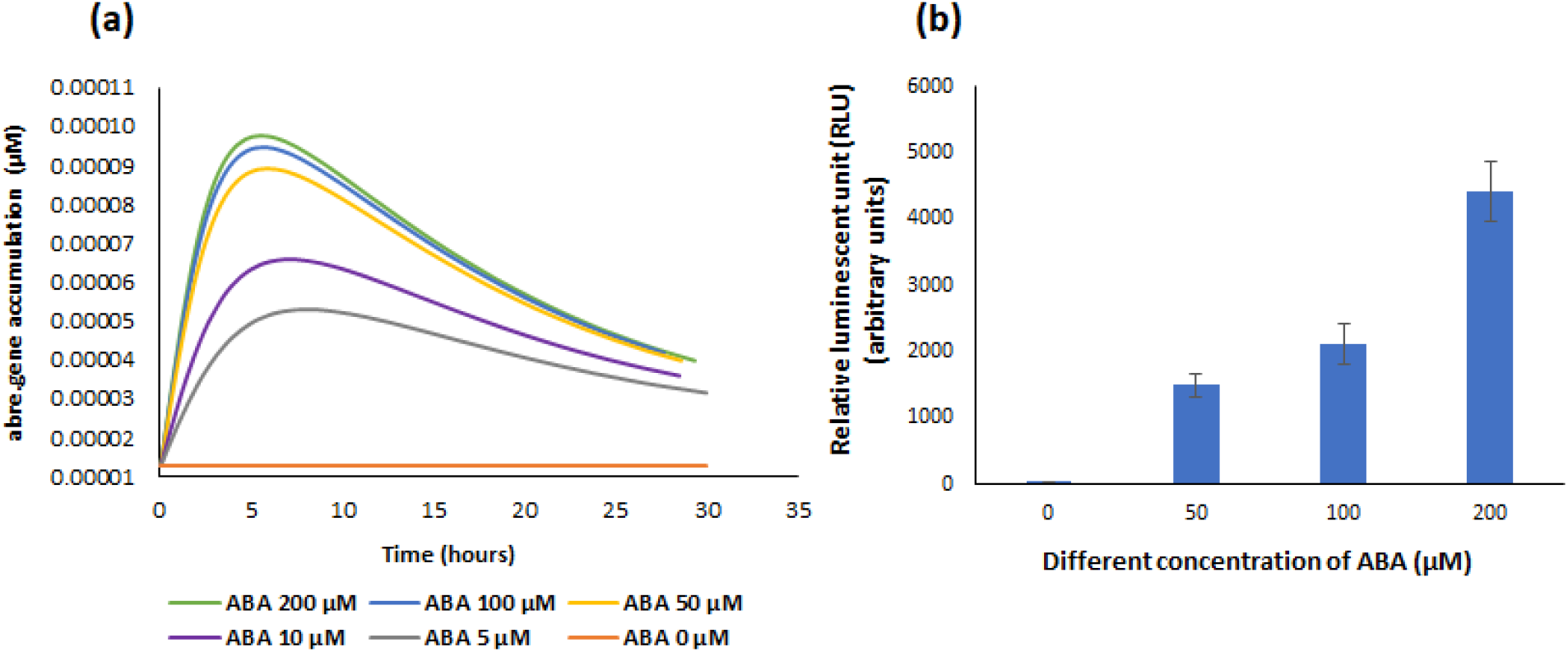
ABRE-promoter activity increases with a function of ABA concentration in the model as it is observed in actual plants. (**a)** Model output of the variable abre.gene with different values of the variable ABA. (**b**) Relative luminescence unit in 25-day-old *RD29A::LUC* plants was determined at 5 hours after spraying different concentrations of ABA. The bars represent the mean relative luminescence of three replicates with error bars representing standard error from the mean (15 seedlings).

We also validated changes of the variable abre.gene in gene-knockout simulations. Namely, we simulated expression of a gene from the ABRE promoter in gene null-mutations of *pyr, pp2c, snrk2*, and *abf*, which were previously studied (Fujita *et al*., 2009; Rubio *et al*., 2009; Nishimura *et al*., 2010; Yoshida *et al*., 2015). We simulated knockout mutations by setting the translation rate constant (kf2) to zero for the variable PYR, PP2C, SnRK2, and ABF. In addition, we also set the translation rates of the feedback regulations kf18 and kf19 to zero for ABF and PP2Cs, respectively. The mimicked null-mutant in *pyr*, *snrk2*, and *abf*, all showed reduced levels of the variable abre.gene while the mimicked null-mutant in *pp2c* showed elevated levels (Table **2**).

**Table 2.** Mutant simulations show similar output to actual mutated plants with respect to the ABRE promoter activity. Mutant simulations were made on the model with the variable ABA set at 100 μM. Highest concentration of the variable abre.gene at each of the simulation was recorded. Relative expression of the *RD29A* gene in actual plants was curated from previously published literatures.

| Variable set to 0 in the model | Highest abre.gene concentration in the model ($\mu\text{M}$) | Knockout genes in actual plants | *RD29A* gene expression in the knockout plants exposed to ABA | Reference |
| --- | --- | --- | --- | --- |
| None | 0.000089 | None (wild type) | transient | (Song *et al.*, 2016) |
| PPC2 | 0.011166 | *pp2ca/hai1* | constitutive and high | (Antoni *et al.*, 2012) |
| PYR | 0.000008 | *pyr1/pyl1/pyl2/pyl4* | impaired | (Park *et al.*, 2009) |
| SnRK2 | 0.000000 | *snrk2.2/ snrk2.3 snrk2.6* | impaired | (Thalmann *et al.*, 2016) |
| ABF | 0.000000 | *areb1/areb2/abf3* | impaired | (Thalmann *et al.*, 2016) |

Experimental data in actual plants shows that *pyr* null-mutants are impaired in ABA-induced gene expression (Park *et al*., 2009; Nishimura *et al*., 2010; Gonzalez-Guzman *et al*., 2012). Similarly, experimental data on *snrk2.2/ snrk2.3/ snrk2.6* triple knockout mutants showed that the expression of ABA-induced genes was impaired (Fujii & Zhu, 2009; Fujita *et al*., 2009; Thalmann *et al*., 2016). Triple *areb/abf* mutants were found to have reduced ABA-induced gene expression (Yoshida *et al*., 2015; Thalmann *et al*., 2016). On the other hand, null-mutants of *pp2cs* in actual plants show a higher and constitutive ABA response (Rubio *et al*., 2009; Antoni *et al*., 2012). Based on the two validations described above, we concluded that the model constructed, and parameters implemented in the model are approximated to actual plants.

### Model simulation and actual plants agree with respect to the activity of ABRE promoter in a condition where PP2C phosphatase activity is inhibited

With the validated model, we examined a relationship between the phosphatase activity of PP2C and the activity of the ABRE promoter, which was not examined before. First, we simulated expression kinetics of the ABA induced gene in which the phosphatase activity of PP2C was decreased. Namely, we decreased the catalytic rate constant of PP2C (kf14). We changed the value from the original 0.924 s^−1^ (Xie *et al*., 2012) to 10^−5^ s^−1^, progressively, and tracked changes of the variable abre.gene for the first 24 hours after changing the variable ABA from 0 to 100 μM (Fig. **4a**).

**Figure 4.**
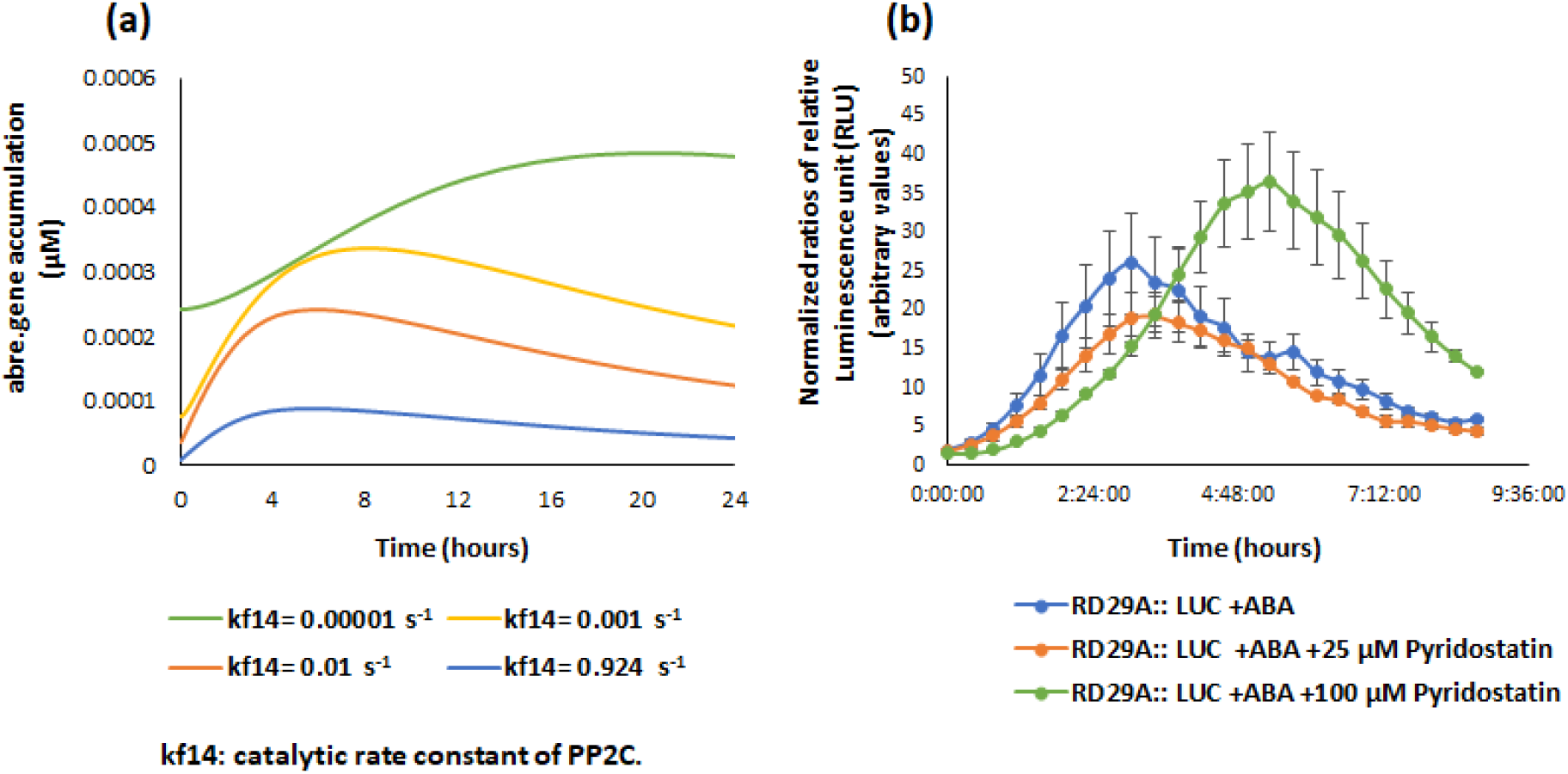
Model simulation and actual plants agree with respect to the activity of ABRE promoter in a condition where PP2C phosphatase activity is inhibited. (**a)** Model simulation for changes in the variable abre.gene. The parameter in catalytic rate constant of PP2C (kf14) is progressively reduced from 0.924 s^−1^ to 10^−5^ s^−1^. Notice the levels of the variable abre.gene increased as the parameter value was reduced. At the same time, the time when the variable abre.gene reached the maximum, was delayed. (**b**) Changes of luminescence in the *RD29A::LUC* transgenic plants. The plants were exposed to pyridostatin, an inhibitor of PP2C phosphatase. The *RD29A::LUC* plants were treated with 100 μM ABA, 100 μM ABA + 25 μM pyridostatin, or 100 μM ABA + 100 μM pyridostatin. Luminescence values were normalized against control (DMSO + 25 μM or 100 μM pyridostatin). Data shown is means of three independent replicates with error bars derived from standard error from the mean. Notice the levels of normalized luminescence intensity was increased and the peak time point was delayed on addition of 100 μM pyridostatin.

On reduction of catalytic rate constant, the variable abre.gene increases, and the peak time point is delayed (Fig. **4a**). Based on the prediction, we hypothesized that inhibition of the PP2C phosphatase activity would increase expression of the ABA induced gene and delay its peak time. To examine the hypothesis, we conducted an experiment with the *RD29A::LUC* transgenic plants and pyridostatin hydrochloride, a recently identified chemical inhibitor that is specific for the PP2C phosphatase activity against SnRK2 (Janicki *et al*., 2020). On addition of 100 μM but not 25 μM pyridostation hydrochloride, an increase in luminescence as well as a delay of the peak time was observed, indicating inhibitor-concentration dependent changes (Fig. **4b**). We also examined the *CAMV35S::LUC* transgenic plants in which a constitutive promoter from a Cauliflower Mosaic Virus drives the expression luciferase (Rosin *et al*., 2008). We observed no significant difference between the plants, in which pyridostation hydrochloride was added or not added, in peak time and luminescence (Fig. **S1**). This confirmed that the change in luminescence kinetics was not due to the alteration of luciferase enzymatic activity, but due to the differential activity of the ABRE promoter. Based on these model predictions and biological experiments, we concluded that inhibition of the PP2C phosphatase activity would increase the ABRE promoter activity and delay its peak time.

### A new hypothesis: ABA downregulates a translation rate of PP2C to increase the ABRE prompter activity

To understand important parameters in the ABA signaling pathway with respect to the ABRE promoter activity, we conducted a sensitive analysis of key parameters against the variable abre.gene in the model.

The analysis found that while most of the selected parameters are equally sensitive, parameters related to ABA and PYR binding were least sensitive. The parameter related to translation of feedbacked PP2Cs, which was optimized in this study to curve-fit the kinetics of the variable abre.gene, had the highest sensitivity (Fig. **5**).

**Figure 5.**
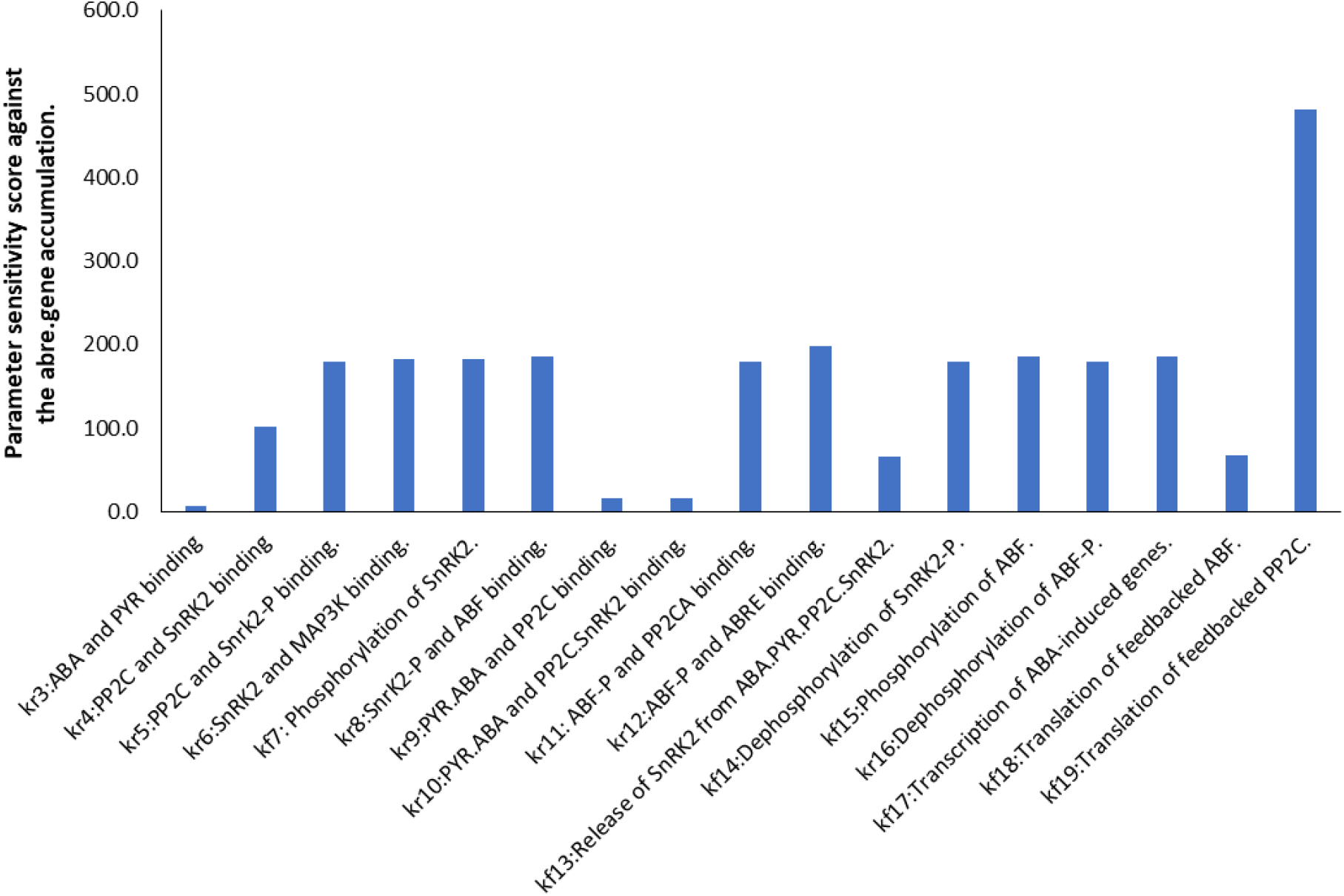
Sensitivity analysis identified the parameter of translation rate constant in feed backed PP2Cs is the most sensitive to the kinetics of the variable abre.gene. A sensitivity analysis was conducted against the variable abre.gene using the calculate sensitivity function in the model analyzer in SimBiology.

To determine how the translation rate constant of PP2Cs affects the ABRE promoter activity, we changed the PP2C translation rate (kf19) and tracked the resulting kinetics of the variable abre.gene. We found that the PP2C translation rate (kf19) affects not only the maximum of variable abre.gene but also the peak time when the highest value of the variable abre.gene is achieved (Fig. **6**). These dynamics are similar to the changes of the parameter in the PP2C enzymatic activity (kf14; Fig. **4a**).

**Figure 6.**
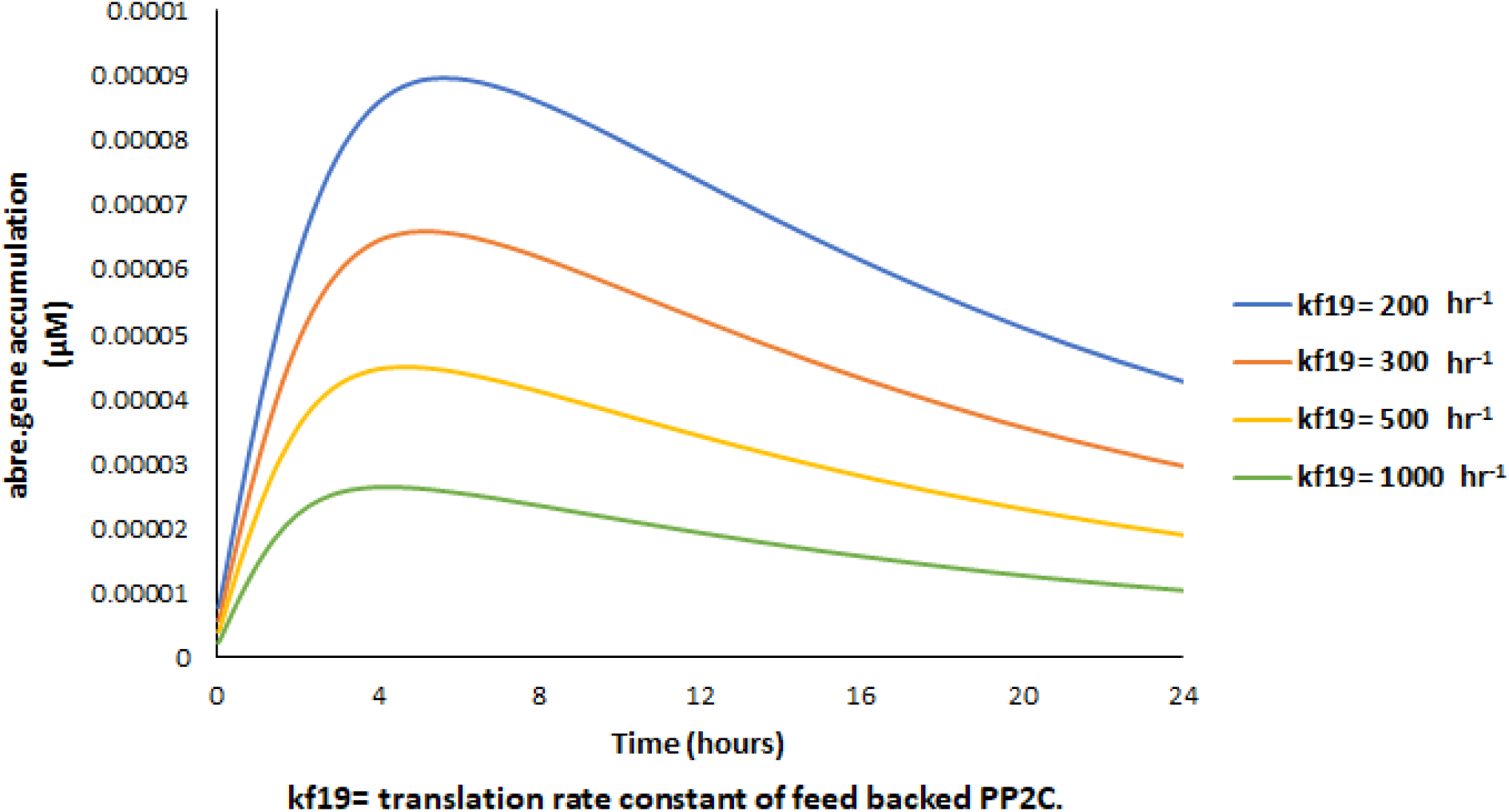
Increase of the translation rate constant of PP2C reduces the variable abre.gene but expedites the peak time. The parameter kf19 (translation rate constant of feed backed PP2C) was changed from the original 200 hr^−1^ to 300, 500, and 1000 hr^−1^. Notice the level and the peak time point of the variable abre.genes changed with a function of translation rate constant.

Learning that the kinetics of the variable abre.gene is largely affected by the translation rate of the feedbacked PP2Cs in the model, we wondered whether the translation rate is affected by ABA in actual plants. To this end, we searched literature that studied changes of the translation rate. We found that while direct measurement of the translation rate in eukaryotic cells has been conducted only in yeast and animal cells (Schwanhäusser *et al*., 2011; Weinberg *et al*., 2016), indirect measurement has been conducted in plants as well (Fujita *et al*., 2019).

In the indirect measurement, using ribosomal profiling, a ratio of ribosome-protected mRNA fragments over total mRNA extracted from cells are measured at a given time point. In theory, a higher ratio of ribosome-protected mRNA over total mRNA indicates higher translation rate at a given time point. We found in a previously conducted study with a DNA microarray that translation rates in all PP2Cs involved in the ABA signaling pathway (namely ABI1, ABI2, HAB1, PP2CA) are downregulated due to dehydration (Table **3**) (Kawaguchi *et al*., 2004). This suggests that the translation rate in PP2Cs may indeed be downregulated by ABA. Because a microarray used in the study does not contain a completed set of gene probes, change in translation rate of ABFs involved in the ABA signaling pathway (namely ABF2, ABF3, and ABF4) is not conclusive. On the other hand, a study with a deep RNA-sequencing technology, in which all extracted mRNAs are measured by sequenced frequency, showed that the translation rates of ABFs involved in the ABA signaling pathway (ABF2, ABF3, and ABF4) are all up-regulated while that of the PP2Cs (data for ABI2 is not available) are little changed upon exposure of exogenously added TOR inhibitor (Scarpin *et al*., 2020) (Table **3**). The study concluded that the plant TOR specifically controls the translation of a set of mRNAs that possesses 5’ oligopyrimidine tract motifs (5’TOPs), which results in alteration of translation in other genes as well.

**Table 3.** Changes of translation rate in PP2Cs and ABFs identified in the previously published data.

| mRNA species. | Relative changes in relative translation rate with dehydration, compared to a control condition (Kawaguchi *et al.*, 2004). | Relative changes in relative translation rate with TOR inhibition, compared to a control condition (Scarpin *et al.*, 2020). |
| --- | --- | --- |
| ABI1 | 0.92 | 1 |
| ABI2 | 0.95 | Data not available |
| HAB1 | 0.80 | 0.92 |
| PP2CA | 0.98 | 1.13 |
| ABF2 | Data not available | 1.39 |
| ABF3 | 0.97 | 1.32 |
| ABF4 | Data not available | 1.15 |

Based on the sensitive analysis on our model and the two previous studies described above, we hypothesized that ABA downregulates a translation rate of PP2C to increase the ABRE prompter activity.

### Combinational exposure of ABA and TOR inhibitor upregulates activity of the ABRE promoter

We further hypothesized that the combinational exposure of ABA and TOR inhibitor up-regulates activity of the ABRE promoter. The rationale is as follows. First, upon ABA exposure, transcription of PP2Cs and ABFs are both upregulated due to the feedback regulation (Wang *et al*., 2019). Secondly, the translation rate of PP2Cs is down regulated by a yet unknown mechanism (Kawaguchi *et al*., 2004), resulting in diminishing the effect of up-regulation of the transcription of PP2Cs. Thirdly, by exposing a TOR inhibitor, translation rate of ABFs is increased while that of PP2Cs is not changed (Scarpin *et al*., 2020). We assumed the increase of the ABF translation occurs independent from the role of TOR in suppression of PYR-ABA binding activity (Wang *et al*., 2018). As a result, by exposing ABA and a TOR inhibitor, the activity of the ABRE promoter increases, compared to when only ABA is exposed to plants.

To examine the hypothesis, we analyzed the ABRE promoter activity in the *RD29A::LUC* transgenic plants. As a control, we analyzed the *CAMV35S::LUC* transgenic plants. We exposed the plants to ABA only and ABA and rapamycin, the TOR inhibitor (Xiong & Sheen, 2012). When the plants were exposed to ABA alone, luciferase intensity was increased as expected (Fig. **7**).

**Figure 7.**
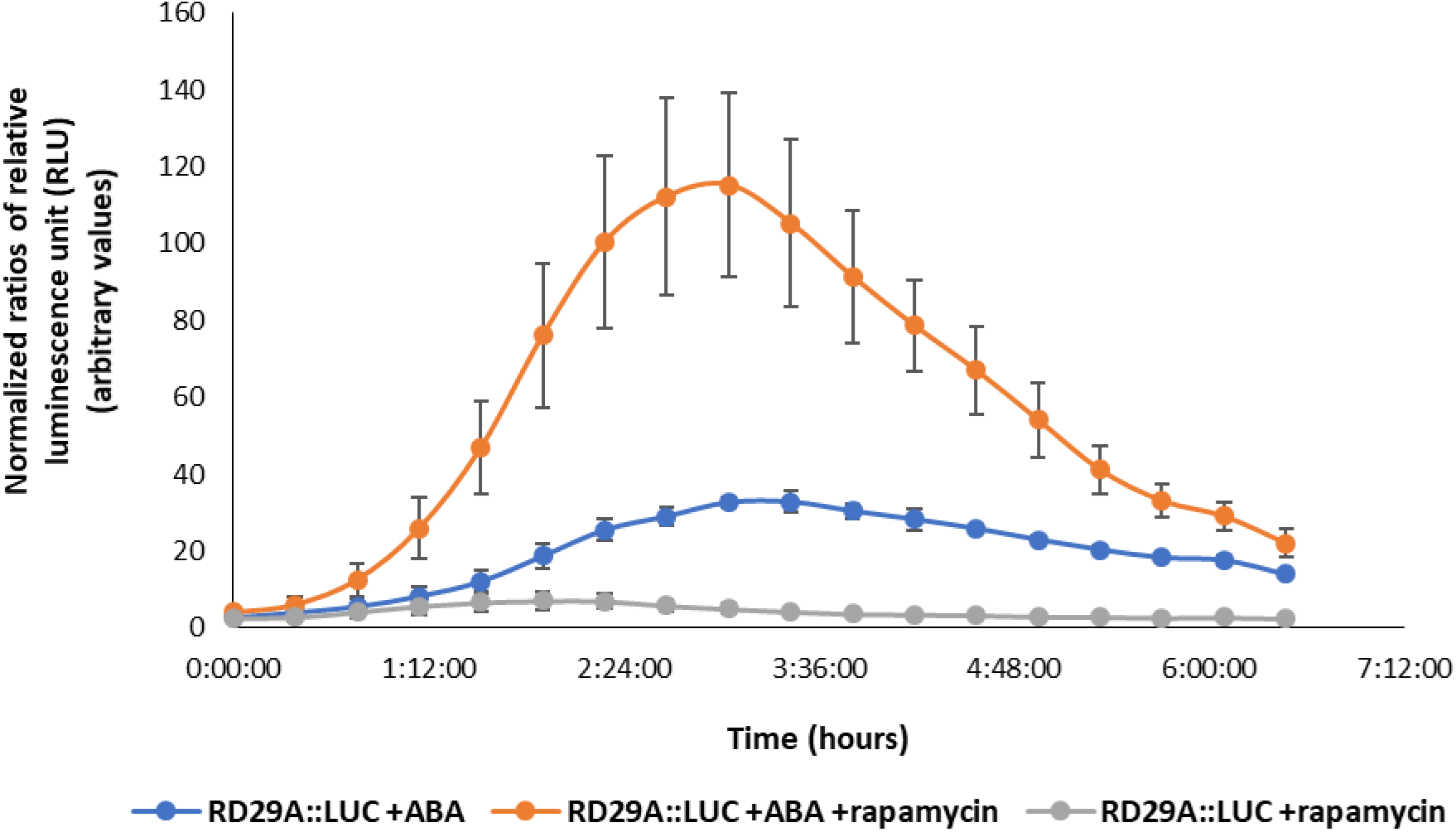
Combinational exposure of ABA and rapamycin increases the ABRE promoter activity. Normalized luminescence in the *RD29A::LUC* transgenic plants are shown. The plants were exposed to 200 μM ABA alone or 200 μM ABA + 10 μM rapamycin or 10 μM rapamycin only. Luminescence values were normalized against control (DMSO only). Data shown is means of three independent replicates with error bars derived from standard error from the mean.

When the plants were exposed to both rapamycin and ABA, the luciferase intensity was about 4-fold higher than that when plants were exposed to ABA alone at the maximum. When the *RD29A::LUC* transgenic plants were exposed to rapamycin alone, luciferase activity was little altered (Fig. **7**). When the *CAMV35S::LUC* transgenic plants were examined with the identical conditions, no significant difference was observed among the different exposures (Fig. **S2**). This result supported our hypothesis that combinational exposure of TOR inhibitor and ABA up-regulates activity of the ABRE promoter.

## Discussion

Here we presented a model of the ABA signaling pathway describing the activation of ABF and resulting activation of the ABRE promoter (Fig. **1**). The model was built with fixed parameter values of protein-protein interactions and enzymatic kinetics that were obtained by *in vitro* experiments from the literature. The model suggests that the feedback regulation of PP2C and ABF allows the transient upregulation of the ABRE promoter. Without the feedback, the model predicts that ABRE expression activity would be logarithmic and not show the transient increase (Fig. **2e**). Based on the model prediction, we hypothesized that inhibition of the PP2C phosphatase activity on SnRK2 would increase expression of the ABA induced gene and delay its peak time. The hypothesis was supported by biological experimentation using transgenic Arabidopsis plants (Fig. **4b**). The model also predicted that the translation rate for PP2C in the feedback regulation is the most sensitive parameter for activation of the ABRE promoter while parameters related to ABA and PYR binding were least sensitive (Fig. **5**). The reason parameters related to ABA and PYR binding were least sensitive is evident because we assume extremely high concentration of ABA (100 μM) is exposed to plants, while a production of endogenous ABA during abiotic stress would be in a nM range (Dubas *et al*., 2013). We found out that a high value of the translation rate not only reduces the ABRE promoter activity but also expedites the time point when the promoter activity reaches the maximum (Fig. **6**). This suggested that the translation rate of PP2C would be one of the most important factors that determine the kinetics of the ABRE promoter activity. In the past, accumulation of mRNA and post-translational modification of proteins are thought to define activity of the ABRE promoter (Nordin *et al*., 1993; Joo *et al*., 2021). However, our model and biological experimental data suggest that changes in translation rates would also largely determine the activity of the ABRE promoter (Fig. **7**). Our literature search found out that the translation rate of PP2Cs is downregulated during dehydration (Table **3**). This suggests that activity of the ABRE promoter would be regulated by not only upregulation of the gene expression but also downregulation of the protein translation on PP2Cs.

We are aware that not only translation rate but also degradation rate of proteins, which are not investigated in this study, are important in the ABA signaling pathway (Wu *et al*., 2016; Ali *et al*., 2019). Hence, changes of protein degradation rate by ABA must be quantitatively analyzed to conclude the role of translation rate in the ABA signaling pathway. We are also aware that ABFs are not the only transcription factors that bind to the ABRE promoter (Song *et al*., 2016). Hence, the activity of the ABRE promoter does not depend only on ABF activation in actual plants, whereas in the model we consider the activity of ABF only. To fully understand kinetics of the ABRE promoter activity in actual plants, further expansion of the model to include other transcription factors is required. Furthermore, quantitative predictions in the current model somewhat disagrees with real plant data. For instance, when an ABA-concentration dependent response of the ABRE promoter was determined, the response range was narrower in the model than in actual plants (Fig. **3**). Optimization of parameter values fixed in this study or the expansion to include other factors driving the ABRE promoter may be required to improve model performance.

Nevertheless, our model successfully builds off existing work to represent the relationship between the ABA signaling pathway and ABRE gene expression. As demonstrated here, the model is useful to generate novel hypotheses. The model suggests new avenues of experimental inquiry. In particular, our analysis proposes that investigating alteration of translation rates in proteins, such as PP2Cs, is the next frontier in the research field of ABA signaling pathway and downstream promoter activity.

## Acknowledgements

This study is, in a part, supported by Economic Development Assistantships from Louisiana State.

## Author Contribution

Conceptualization and methodology, N.K. Validation, R.N. and R.D. Experiments, R.N. Formal analysis, R.N. and N.K. Writing—original draft preparation, R.N. and N.K Writing—review and editing, R.N. R.D. and N.K. Funding acquisition, N.K. All authors have read and agreed to the published version of the manuscript.

## Data Availability

.sbproj file (MATLAB SimBiology Project File) that includes a model diagram, ODE equations, initial values, parameters, simulations for Figures 2, 3, 4, 5, 6, and Table 2 are available as supplement files.

## References

Albert R, Acharya BR, Jeon BW, Zañudo JGT, Zhu M, Osman K, Assmann SM. 2017. A new discrete dynamic model of ABA-induced stomatal closure predicts key feedback loops. PLOS Biology 15: e2003451.

Aldridge BB, Burke JM, Lauffenburger DA, Sorger PK. 2006. Physicochemical modelling of cell signalling pathways. Nature Cell Biology 8.

Ali A, Kim JK, Jan M, Khan HA, Khan IU, Shen M, Park J, Lim CJ, Hussain S, Baek D, et al. 2019. Rheostatic Control of ABA Signaling through HOS15-Mediated OST1 Degradation. Molecular Plant 12: 1447–1462.

Antoni R, Gonzalez-Guzman M, Rodriguez L, Rodrigues A, Pizzio GA, Rodriguez PL. 2012. Selective Inhibition of Clade A Phosphatases Type 2C by PYR/PYL/RCAR Abscisic Acid Receptors. Plant Physiology 158: 970–980.

Bar-Even A, Noor E, Savir Y, Liebermeister W, Davidi D, Tawfik DS, Milo R. 2011. The Moderately Efficient Enzyme: Evolutionary and Physicochemical Trends Shaping Enzyme Parameters. Biochemistry 50: 4402–4410.

Basu S, Ramegowda V, Kumar A, Pereira A. 2016. Plant adaptation to drought stress. F1000Research 5: F1000 Faculty Rev-1554.

Belin C, de Franco P-O, Bourbousse C, Chaignepain S, Schmitter J-M, Vavasseur A, Giraudat J, Barbier-Brygoo H, Thomine S. 2006. Identification of features regulating OST1 kinase activity and OST1 function in guard cells. Plant Physiology 141: 1316–1327.

Choi H, Hong J, Ha J, Kang J, Kim SY. 2000. ABFs, a Family of ABA-responsive Element Binding Factors *. Journal of Biological Chemistry 275: 1723–1730.

Dubas E, Janowiak F, Krzewska M, Hura T, Żur I. 2013. Endogenous ABA concentration and cytoplasmic membrane fluidity in microspores of oilseed rape (Brassica napus L.) genotypes differing in responsiveness to androgenesis induction. Plant Cell Reports 32: 1465–1475.

Dupeux F, Santiago J, Betz K, Twycross J, Park S-Y, Rodriguez L, Gonzalez-Guzman M, Jensen MR, Krasnogor N, Blackledge M, et al. 2011. A thermodynamic switch modulates abscisic acid receptor sensitivity. The EMBO Journal 30: 4171–4184.

Forzani C, Duarte GT, Van Leene J, Clément G, Huguet S, Paysant-Le-Roux C, Mercier R, De Jaeger G, Leprince A-S, Meyer C. 2019. Mutations of the AtYAK1 Kinase Suppress TOR Deficiency in Arabidopsis. Cell Reports 27: 3696–3708.e5.

Fujii H, Chinnusamy V, Rodrigues A, Rubio S, Antoni R, Park S-Y, Cutler SR, Sheen J, Rodriguez PL, Zhu J-K. 2009. In vitro Reconstitution of an ABA Signaling Pathway. Nature 462: 660–664.

Fujii H, Zhu J-K. 2009. Arabidopsis mutant deficient in 3 abscisic acid-activated protein kinases reveals critical roles in growth, reproduction, and stress. Proceedings of the National Academy of Sciences 106: 8380–8385.

Fujita T, Kurihara Y, Iwasaki S. 2019. The Plant Translatome Surveyed by Ribosome Profiling. Plant & Cell Physiology 60: 1917–1926.

Fujita Y, Nakashima K, Yoshida T, Katagiri T, Kidokoro S, Kanamori N, Umezawa T, Fujita M, Maruyama K, Ishiyama K, et al. 2009. Three SnRK2 Protein Kinases are the Main Positive Regulators of Abscisic Acid Signaling in Response to Water Stress in Arabidopsis. Plant and Cell Physiology 50: 2123–2132.

Gampala SSL, Hagenbeek D, Rock CD. 2001. Functional Interactions of Lanthanum and Phospholipase D with the Abscisic Acid Signaling Effectors VP1 and ABI1-1 in Rice Protoplasts. Journal of Biological Chemistry 276: 9855–9860.

Geertz M, Shore D, Maerkl SJ. 2012. Massively parallel measurements of molecular interaction kinetics on a microfluidic platform. Proceedings of the National Academy of Sciences 109: 16540–16545.

Geiger D, Scherzer S, Mumm P, Stange A, Marten I, Bauer H, Ache P, Matschi S, Liese A, Al-Rasheid KAS, et al. 2009. Activity of guard cell anion channel SLAC1 is controlled by drought-stress signaling kinase-phosphatase pair. Proceedings of the National Academy of Sciences of the United States of America 106: 21425–21430.

Ghose R. 2019. Nature of the Pre-Chemistry Ensemble in Mitogen-Activated Protein Kinases. Journal of Molecular Biology 431: 145–157.

Gonzalez-Guzman M, Pizzio GA, Antoni R, Vera-Sirera F, Merilo E, Bassel GW, Fernández MA, Holdsworth MJ, Perez-Amador MA, Kollist H, et al. 2012. Arabidopsis PYR/PYL/RCAR Receptors Play a Major Role in Quantitative Regulation of Stomatal Aperture and Transcriptional Response to Abscisic Acid. The Plant Cell 24: 2483–2496.

Gosti F, Beaudoin N, Serizet C, Webb AAR, Vartanian N, Giraudat J. 1999. ABI1 Protein Phosphatase 2C Is a Negative Regulator of Abscisic Acid Signaling. The Plant Cell 11: 1897–1909.

Hausser J, Mayo A, Keren L, Alon U. 2019. Central dogma rates and the trade-off between precision and economy in gene expression. Nature Communications 10: 68.

Hoops S, Hontecillas R, Abedi V, Leber A, Philipson C, Carbo A, Bassaganya-Riera J. 2016. Chapter 5 - Ordinary Differential Equations (ODEs) Based Modeling. In: Bassaganya-Riera J, ed. Computational Immunology. Academic Press, 63–78.

Ikegami K, Okamoto M, Seo M, Koshiba T. 2008. Activation of abscisic acid biosynthesis in the leaves of Arabidopsis thaliana in response to water deficit. Journal of Plant Research 122: 235.

Janes KA, Yaffe MB. 2006. Data-driven modelling of signal-transduction networks. Nature Reviews Molecular Cell Biology 7.

Janicki M, Marczak M, Cieśla A, Ludwików A. 2020. Identification of Novel Inhibitors of a Plant Group A Protein Phosphatase Type 2C Using a Combined In Silico and Biochemical Approach. Frontiers in Plant Science 11: 1416.

Joo H, Baek W, Lim CW, Lee SC. 2021. Post-translational Modifications of bZIP Transcription Factors in Abscisic Acid Signaling and Drought Responses. Current Genomics 22: 4–15.

Kawaguchi R, Girke T, Bray EA, Bailey-Serres J. 2004. Differential mRNA translation contributes to gene regulation under non-stress and dehydration stress conditions in Arabidopsis thaliana. The Plant Journal: For Cell and Molecular Biology 38: 823–839.

Kim D, Ntui VO, Xiong L. 2016. Arabidopsis YAK1 regulates abscisic acid response and drought resistance. FEBS letters 590: 2201–2209.

Kumar S, Sachdeva S, Bhat KV, Vats S. 2018. Plant Responses to Drought Stress: Physiological, Biochemical and Molecular Basis. In: Vats S, ed. Biotic and Abiotic Stress Tolerance in Plants. Singapore: Springer, 1–25.

Lee SY, Boon NJ, Webb AAR, Tanaka RJ. 2016. Synergistic Activation of RD29A Via Integration of Salinity Stress and Abscisic Acid in Arabidopsis thaliana. Plant and Cell Physiology 57: 2147–2160.

Lee SC, Lan W, Buchanan BB, Luan S. 2009. A protein kinase-phosphatase pair interacts with an ion channel to regulate ABA signaling in plant guard cells. Proceedings of the National Academy of Sciences of the United States of America 106: 21419–21424.

Li S, Assmann SM, Albert R. 2006. Predicting Essential Components of Signal Transduction Networks: A Dynamic Model of Guard Cell Abscisic Acid Signaling. PLOS Biology 4: e312.

Lynch T, Erickson BJ, Finkelstein RR. 2012. Direct interactions of ABA-insensitive(ABI)-clade protein phosphatase(PP)2Cs with calcium-dependent protein kinases and ABA response element-binding bZIPs may contribute to turning off ABA response. Plant Molecular Biology 80: 647–658.

Ma Y, Szostkiewicz I, Korte A, Moes D, Yang Y, Christmann A, Grill E. 2009. Regulators of PP2C Phosphatase Activity Function as Abscisic Acid Sensors. Science 324: 1064–1068.

Maheshwari P, Assmann SM, Albert R. 2020. A Guard Cell Abscisic Acid (ABA) Network Model That Captures the Stomatal Resting State. Frontiers in Physiology 0.

Maheshwari P, Du H, Sheen J, Assmann SM, Albert R. 2019. Model-driven discovery of calcium-related protein-phosphatase inhibition in plant guard cell signaling. PLOS Computational Biology 15: e1007429.

Melcher K, Ng L-M, Zhou XE, Soon F-F, Xu Y, Suino-Powell KM, Park S-Y, Weiner JJ, Fujii H, Chinnusamy V, et al. 2009. A Gate-Latch-Lock Mechanism for Hormone Signaling by Abscisic Acid Receptors. Nature 462: 602–608.

Merlot S, Gosti F, Guerrier D, Vavasseur A, Giraudat J. 2001. The ABI1 and ABI2 protein phosphatases 2C act in a negative feedback regulatory loop of the abscisic acid signalling pathway. The Plant Journal: For Cell and Molecular Biology 25: 295–303.

Milo R, Phillips R. 2015. Cell Biology by the numbers. New York: Garland Science.

Murashige T, Skoog F. 1962. A Revised Medium for Rapid Growth and Bio Assays with Tobacco Tissue Cultures. Physiologia Plantarum 15: 473–497.

Nishimura N, Hitomi K, Arvai AS, Rambo RP, Hitomi C, Cutler SR, Schroeder JI, Getzoff ED. 2009. Structural Mechanism of Abscisic Acid Binding and Signaling by Dimeric PYR1. Science (New York, N.Y.) 326: 1373–1379.

Nishimura N, Sarkeshik A, Nito K, Park S-Y, Wang A, Carvalho PC, Lee S, Caddell DF, Cutler SR, Chory J, et al. 2010. PYR/PYL/RCAR family members are major in-vivo ABI1 protein phosphatase 2C-interacting proteins in Arabidopsis. The Plant Journal: For Cell and Molecular Biology 61: 290–299.

Nishimura N, Yoshida T, Kitahata N, Asami T, Shinozaki K, Hirayama T. 2007. ABA-Hypersensitive Germination1 encodes a protein phosphatase 2C, an essential component of abscisic acid signaling in Arabidopsis seed. The Plant Journal: For Cell and Molecular Biology 50: 935–949.

Nordin K, Vahala T, Palva ET. 1993. Differential expression of two related, low-temperature-induced genes in Arabidopsis thaliana (L.) Heynh. Plant Molecular Biology 21: 641–653.

Norval LW, Krämer SD, Gao M, Herz T, Li J, Rath C, Wöhrle J, Günther S, Roth G. 2019. KOFFI and Anabel 2.0—a new binding kinetics database and its integration in an open-source binding analysis software. Database 2019.

Pan C, Tang J, Xu Y, Xiao P, Liu H, Wang H, Wang W, Meng F, Yu X, Sun J. 2015. The catalytic role of the M2 metal ion in PP2Cα. Scientific Reports 5.

Park S-Y, Fung P, Nishimura N, Jensen DR, Fujii H, Zhao Y, Lumba S, Santiago J, Rodrigues A, Chow TF, et al. 2009. Abscisic acid inhibits PP2Cs via the PYR/PYL family of ABA-binding START proteins. Science (New York, N.Y.) 324: 1068–1071.

Philips RM & R. Cell Biology by the Numbers.

Poolman MG, Assmus HE, Fell DA. 2004. Applications of metabolic modelling to plant metabolism. Journal of Experimental Botany 55: 1177–1186.

Rodriguez PL, Leube MP, Grill E. 1998. Molecular cloning in Arabidopsis thaliana of a new protein phosphatase 2C (PP2C) with homology to ABI1 and ABI2. Plant Molecular Biology 38: 879–883.

Rosin FM, Watanabe N, Cacas J-L, Kato N, Arroyo JM, Fang Y, May B, Vaughn M, Simorowski J, Ramu U, et al. 2008. Genome-wide transposon tagging reveals location-dependent effects on transcription and chromatin organization in Arabidopsis. The Plant Journal: For Cell and Molecular Biology 55: 514–525.

Rubio S, Rodrigues A, Saez A, Dizon MB, Galle A, Kim T-H, Santiago J, Flexas J, Schroeder JI, Rodriguez PL. 2009. Triple Loss of Function of Protein Phosphatases Type 2C Leads to Partial Constitutive Response to Endogenous Abscisic Acid. Plant Physiology 150: 1345–1355.

Saez A, Apostolova N, Gonzalez-Guzman M, Gonzalez-Garcia MP, Nicolas C, Lorenzo O, Rodriguez PL. 2004. Gain-of-function and loss-of-function phenotypes of the protein phosphatase 2C HAB1 reveal its role as a negative regulator of abscisic acid signalling. The Plant Journal: For Cell and Molecular Biology 37: 354–369.

Santiago J, Rodrigues A, Saez A, Rubio S, Antoni R, Dupeux F, Park S-Y, Márquez JA, Cutler SR, Rodriguez PL. 2009. Modulation of drought resistance by the abscisic acid receptor PYL5 through inhibition of clade A PP2Cs. The Plant Journal 60: 575–588.

Sauter A, Davies WJ, Hartung W. 2001. The long-distance abscisic acid signal in the droughted plant: the fate of the hormone on its way from root to shoot. Journal of Experimental Botany 52: 1991–1997.

Scarpin MR, Leiboff S, Brunkard JO. 2020. Parallel global profiling of plant TOR dynamics reveals a conserved role for LARP1 in translation (JL Manley, N Sonenberg, and O Meyuhas, Eds.). eLife 9: e58795.

Schroeder JI, Hedrich R, Fernandez JM. 1984. Potassium-selective single channels in guard cell protoplasts of Vicia faba. Nature 312: 361–362.

Schwanhäusser B, Busse D, Li N, Dittmar G, Schuchhardt J, Wolf J, Chen W, Selbach M. 2011. Global quantification of mammalian gene expression control. Nature 473: 337–342.

Singh A, Jha SK, Bagri J, Pandey GK. 2015. ABA Inducible Rice Protein Phosphatase 2C Confers ABA Insensitivity and Abiotic Stress Tolerance in Arabidopsis. PLOS ONE 10: e0125168.

Song L, Huang SC, Wise A, Castanon R, Nery JR, Chen H, Watanabe M, Thomas J, Bar-Joseph Z, Ecker JR. 2016. A transcription factor hierarchy defines an environmental stress response network. Science 354.

Soon F-F, Ng L-M, Zhou XE, West GM, Kovach A, Tan MHE, Suino-Powell KM, He Y, Xu Y, Chalmers MJ, et al. 2012. Molecular Mimicry Regulates ABA Signaling by SnRK2 Kinases and PP2C Phosphatases. Science (New York, N.Y.) 335: 85–88.

Steuer B, Stuhlfauth T, Fock HP. 1988. The efficiency of water use in water stressed plants is increased due to ABA induced stomatal closure. Photosynthesis Research 18: 327–336.

Takahashi F, Kuromori T, Urano K, Yamaguchi-Shinozaki K, Shinozaki K. 2020a. Drought Stress Responses and Resistance in Plants: From Cellular Responses to Long-Distance Intercellular Communication. Frontiers in Plant Science 11: 1407.

Takahashi Y, Zhang J, Hsu P-K, Ceciliato PHO, Zhang L, Dubeaux G, Munemasa S, Ge C, Zhao Y, Hauser F, et al. 2020b. MAP3Kinase-dependent SnRK2-kinase activation is required for abscisic acid signal transduction and rapid osmotic stress response. Nature Communications 11: 12.

Thakar J, Pilione M, Kirimanjeswara G, Harvill ET, Albert R. 2007. Modeling Systems-Level Regulation of Host Immune Responses. PLOS Computational Biology 3: e109.

Thalmann M, Pazmino D, Seung D, Horrer D, Nigro A, Meier T, Kölling K, Pfeifhofer HW, Zeeman SC, Santelia D. 2016. Regulation of Leaf Starch Degradation by Abscisic Acid Is Important for Osmotic Stress Tolerance in Plants[OPEN]. The Plant Cell 28: 1860–1878.

Umezawa T, Sugiyama N, Mizoguchi M, Hayashi S, Myouga F, Yamaguchi-Shinozaki K, Ishihama Y, Hirayama T, Shinozaki K. 2009. Type 2C protein phosphatases directly regulate abscisic acid-activated protein kinases in Arabidopsis. Proceedings of the National Academy of Sciences 106: 17588–17593.

Uno Y, Furihata T, Abe H, Yoshida R, Shinozaki K, Yamaguchi-Shinozaki K. 2000. Arabidopsis basic leucine zipper transcription factors involved in an abscisic acid-dependent signal transduction pathway under drought and high-salinity conditions. Proceedings of the National Academy of Sciences 97: 11632–11637.

Wang X, Guo C, Peng J, Li C, Wan F, Zhang S, Zhou Y, Yan Y, Qi L, Sun K, et al. 2019. ABRE-BINDING FACTORS play a role in the feedback regulation of ABA signaling by mediating rapid ABA induction of ABA co-receptor genes. New Phytologist 221: 341–355.

Wang P, Zhao Y, Li Z, Hsu C-C, Liu X, Fu L, Hou Y-J, Du Y, Xie S, Zhang C, et al. 2018. Reciprocal Regulation of the TOR Kinase and ABA Receptor Balances Plant Growth and Stress Response. Molecular Cell 69: 100–112.e6.

Weinberg DE, Shah P, Eichhorn SW, Hussmann JA, Plotkin JB, Bartel DP. 2016. Improved Ribosome-Footprint and mRNA Measurements Provide Insights into Dynamics and Regulation of Yeast Translation. Cell Reports 14: 1787–1799.

Wu Q, Zhang X, Peirats-Llobet M, Belda-Palazon B, Wang X, Cui S, Yu X, Rodriguez PL, An C. 2016. Ubiquitin Ligases RGLG1 and RGLG5 Regulate Abscisic Acid Signaling by Controlling the Turnover of Phosphatase PP2CA. The Plant Cell 28: 2178–2196.

Xie T, Ren R, Zhang Y, Pang Y, Yan C, Gong X, He Y, Li W, Miao D, Hao Q, et al. 2012. Molecular Mechanism for Inhibition of a Critical Component in the Arabidopsis thaliana Abscisic Acid Signal Transduction Pathways, SnRK2.6, by Protein Phosphatase ABI1. The Journal of Biological Chemistry 287: 794–802.

Xiong Y, Sheen J. 2012. Rapamycin and Glucose-Target of Rapamycin (TOR) Protein Signaling in Plants*. Journal of Biological Chemistry 287: 2836–2842.

Yin P, Fan H, Hao Q, Yuan X, Wu D, Pang Y, Yan C, Li W, Wang J, Yan N. 2009. Structural insights into the mechanism of abscisic acid signaling by PYL proteins. Nature Structural & Molecular Biology 16: 1230–1236.

Yoshida T, Fujita Y, Maruyama K, Mogami J, Todaka D, Shinozaki K, Yamaguchi-Shinozaki K. 2015. Four Arabidopsis AREB/ABF transcription factors function predominantly in gene expression downstream of SnRK2 kinases in abscisic acid signalling in response to osmotic stress. Plant, Cell & Environment 38: 35–49.

Zeevaart JAD, Creelman RA. 1988. Metabolism and Physiology of Abscisic Acid. Annual Review of Plant Physiology and Plant Molecular Biology 39: 439–473.

Zhan X, Wang B, Li H, Liu R, Kalia RK, Zhu J-K, Chinnusamy V. 2012. Arabidopsis proline-rich protein important for development and abiotic stress tolerance is involved in microRNA biogenesis. Proceedings of the National Academy of Sciences 109: 18198–18203.

Zhao J, Zhao L, Zhang M, Zafar SA, Fang J, Li M, Zhang W, Li X. 2017. Arabidopsis E3 Ubiquitin Ligases PUB22 and PUB23 Negatively Regulate Drought Tolerance by Targeting ABA Receptor PYL9 for Degradation. International Journal of Molecular Sciences 18: 1841.

