## Supplementary materials for "A dynamic model of the ABA Signaling pathway with its core components: translation rate of PP2C determines the kinetics of ABA-induced gene expression"

### Supplemental Figures

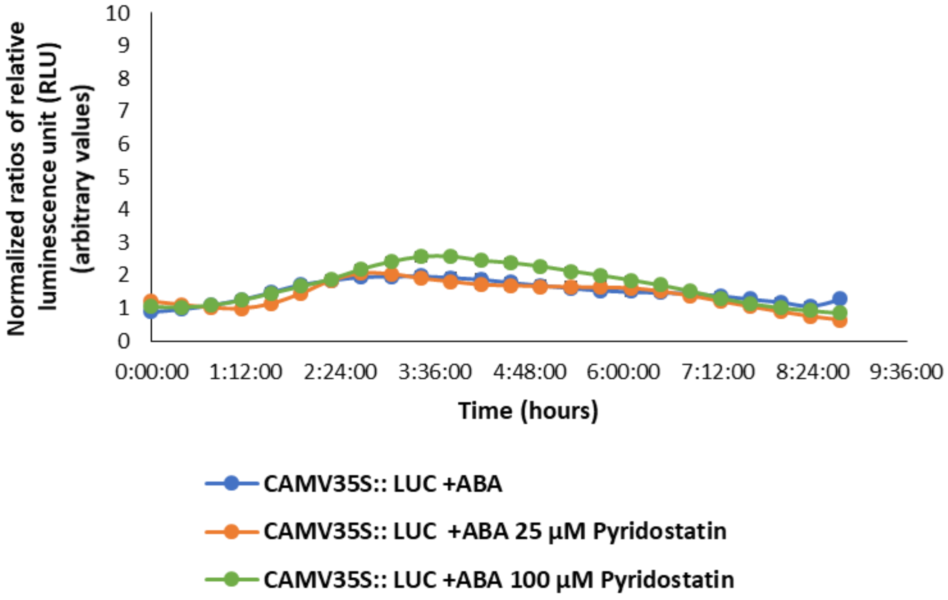


**Figure S1. *CAMV35S::LUC* expression is not significantly different between combinational exposure of ABA and pyridostatin in comparison to ABA only.** *CAMV35S::LUC* plants were exposed to pyridostatin, the inhibitor of PP2C phosphatase against SnRK2. The *CAMV35S::LUC* plants were treated with 100 µM ABA, 100 µM ABA + 25 µM pyridostatin or 100 µM ABA + 100 µM pyridostatin. Luminescence values were normalized against control (DMSO + 25 µM or 100 µM pyridostatin exposure). Data shown is means of three independent replicates with error bars derived from standard error from the mean.

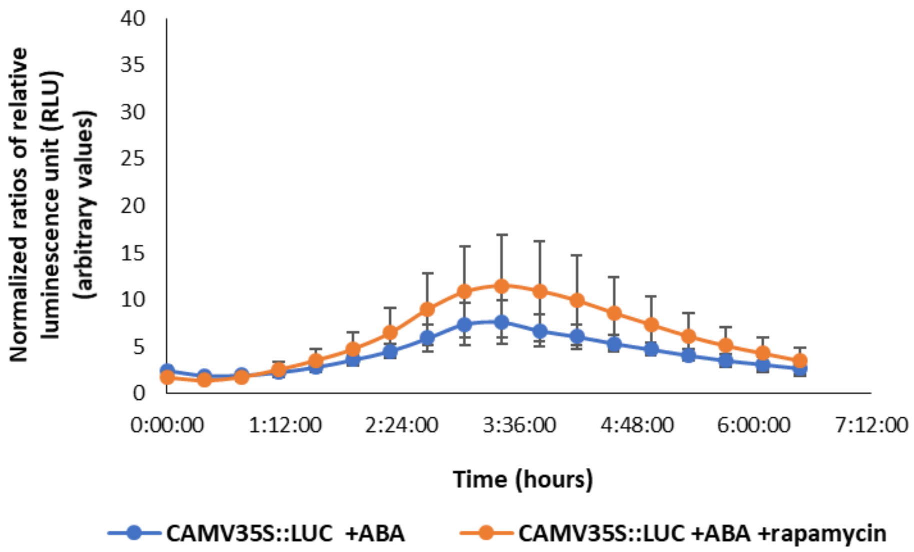


**Figure S2. *CAMV35S::LUC* expression is not significantly different between combinational exposure of ABA and rapamycin in comparison to ABA only.** Normalized luminescence in the *CAMV35S::LUC* transgenic plants are shown. The plants were exposed to 200 µM ABA alone or 200 µM ABA + 10 µM rapamycin. Luminescence values were normalized against control (DMSO only). Data shown is means of three independent replicates with error bars derived from standard error from the mean.

### Supplementary methods

#### Transgenic plant and growth conditions

Transgenic *Arabidopsis thaliana* seeds (CS67900) that carry a *RD29A::LUC* gene expression cassette in the genome and transgenic *Arabidopsis thaliana* seeds (CS25237 or CS25230) that carry a *CAMV35S::LUC* gene expression cassette in the genome were obtained from Arabidopsis Biological Resource Center (ABRC). They were sterilized in 70% ethanol for 1 minute then in 50% bleach, 0.05% triton for 10 minutes. The seeds were then rinsed 6 times with sterilized distilled water, and then plated on ½ Murashige-Skoog (MS) medium (Murashige & Skoog, 1962) containing 0.8% agar. They were then stratified at 4 $^o$ C for 3 days, then placed in a growth chamber under a growth cycle of 16 hours day 8 hours dark with light set at 100 µmol $m^{-2}$ $s^{-1}$ and at 22 $^o$ C.

#### Long term luminescence assay with transgenic *Arabidopsis thaliana*

Twenty-five-day old transgenic *Arabidopsis thaliana* plants on an agar plate were sprayed with 200 µM ABA and for control with the same concentration (v/v) of DMSO for different time periods (0, 5, 8, 12, and 24 hours). Fifteen seedlings were harvested at each time point and frozen immediately in liquid $N_2$ and kept at –80$^o$C until analysis. For analysis, frozen plants were ground using a mortar and pestle to form a powder. The powder was mixed with passive lysis buffer (Promega: Cat. #E1941). The mix was then vortexed and centrifuged at 14,000rpm for 10 minutes. The supernatant (40 µL) was placed in a well on a 96 well plate (Thermo Fischer cat# 267350) and mixed with 100 µL luciferase assay substrate (Promega: Cat. #E151A). Relative Luminescence Unit (RLU) was detected using a Veritas ™ microplate luminometer. Three RLU readings were made for each well and then averaged as data.

#### Short term luminescence assay with transgenic *Arabidopsis thaliana*

One-week-old transgenic *Arabidopsis thaliana* seedlings were placed in wells in a 96-well plate. 100 µL of ½ Murashige-Skoog solution with DMSO, ABA or ABA + rapamycin (Millipore-Sigma cat#53123-88-9) or ABA + pyridostatin hydrochloride (Sigma Aldrich cat# SML2690) was added to each well. Three and one seedlings of *RD29A::LUC* and *CAMV35S::LUC* transgenic plants were placed in each well, respectively. Biological replicates were obtained from 6 wells for each treatment. RLU was detected in a Veritas $^{TM}$ microplate luminometer every 20 min for 8 hours after adding1mM of D-luciferin (ThermoFisher: Cat. #88293) in each well.
